# MATERNAL ZIKA VIRUS (ZIKV) INFECTION FOLLOWING VAGINAL INOCULATION WITH ZIKV-INFECTED SEMEN IN THE TIMED-PREGNANT OLIVE BABOON

**DOI:** 10.1101/2020.01.10.902692

**Authors:** Sunam Gurung, Hugh Nadeau, Marta Maxted, Jamie Peregrine, Darlene Reuter, Abby Norris, Rodney Edwards, Kimberly Hyatt, Krista Singleton, James F. Papin, Dean A. Myers

## Abstract

ZIKV infection is associated with pregnancy loss, fetal microcephaly and other malformations. While *Aedes sp.* of mosquito are the primary vector for ZIKV, sexual transmission of ZIKV is a significant route of infection. ZIKV has been documented in human, mouse and non-human primate (NHP) semen. It is critical to establish NHP models of vertical transfer of ZIKV that recapitulate human ZIKV pathogenesis. We hypothesized that vaginal deposition of ZIKV infected baboon semen would lead to maternal infection and vertical transfer in the olive baboon (*Papio anubis*). Timed pregnant baboons (n=6) were inoculated via vaginal deposition of baboon semen containing 10^6^ ffu ZIKV (n=3, French Polynesian isolate:H/PF/2013, n=3 Puerto Rican isolate:PRVABC59) at mid-gestation (86-95 days gestation [dG]; term 183dG) on day (d) 0 (all dams), and then at 7 day intervals through three weeks. Maternal blood, saliva and cervico-vaginal washes were obtained at select days post-inoculation. Animals were euthanized at 28 days post initial inoculation (dpi; n=5) or 39 dpi (n=1) and maternal/fetal tissues collected. vRNA was quantified by qPCR. Viremia was achieved in 3/3 FP ZIKV infected dams and 2/3 PR ZIKV. ZIKV RNA was detected in cvw (5/6 dams;). ZIKV RNA was detected in lymph nodes, but not ovary, uterus, cervix or vagina in the FP ZIKV dams but was detected in uterus, vagina and lymph nodes. Placenta, amniotic fluid and all fetal tissues were ZIKV RNA negative in the FP infected dams whereas 2/3 PR infected dam placentas were ZIKV RNA positive. We conclude that ZIKV infected semen is a means of ZIKV transmission during pregnancy in primates. The PR isolate appeared more capable of wide spread dissemination to tissues, including placenta compared to the FP strain.

**IMPORTANCE:** Due to its established link to pregnancy loss, microcephaly and other major congenital anomalies, Zika virus (ZIKV) remains a worldwide health threat. Although mosquitoes are the primary means of ZIVK transmission, sexual transmission in human populations is well documented and provides a means for widespread dissemination of the virus. Differences in viremia, tissue distribution, immune responses and pregnancy outcome from sexually transmitted ZIKV compared to the subcutaneous route of infection are needed to better clinically manage ZIKV in pregnancy. Through our previous work, we have developed the olive baboon as a non-human primate model of ZIKV infection that is permissible to ZIKV infection via the subcutaneous route of inoculation and transfer of ZIKV to the fetus in pregnancy. The current study evaluated the course of ZIKV infection after vaginal inoculation of ZIKV in pregnant baboons at mid-gestation using baboon semen as the carrier and comparing two isolates of ZIKV, the French Polynesian isolate first associated with microcephaly and the Puerto Rican isolate, associated with an increased risk of microcephaly observed in the Americas.

## INTRODUCTION

The propagation of Zika virus (ZIKV) represents a worldwide reproductive health crisis given the virus’s geographic distribution and severity of its effect on the developing fetus. At its peak, ZIKV infection was being reported in more than 60 countries (1, 2). While ZIKV infection is usually mild or asymptomatic with typical symptoms that include fever, rash, and conjunctivitis, the impact of ZIKV on the developing fetus can be severe, including microcephaly and associated neurological damage, fetal growth restriction, intrauterine fetal demise, spontaneous abortion as well as other congenital anomalies now termed collectively as Congenital Zika Syndrome (CZS). The impact of ZIKV infection on the fetus led to worldwide interest in the virus following the major outbreak in Brazil in 2015 (3).

While the *Aedes* species of mosquitos are the primary means of ZIKV transmission, sexual transmission of ZIKV is now well documented in the human population (4–8) and a greater incidence of ZIKV in females of a sexually active age (9) suggesting sexual transmission as a causative factor. Sexual transmission in humans was first reported in 2008 (10) and from 2015-2018, 52 cases of sexually transmitted ZIKV were reported in the United States (11). Sexual transmission of ZIKV has been described from a male to pregnant female as well (12, 13). ZIKV RNA and infectious virus have been detected in human semen, considerably longer than detection in blood, saliva and urine (6). ZIKV RNA has been detected in semen at 188 days post-onset of symptoms, while infectious ZIKV has been observed in human semen up to 69 days post-symptoms (8, 14). We recently demonstrated ZIKV RNA in semen in male baboons up to 41 days post infection which was when the study was terminated and persistence of ZIKV RNA in the male reproductive tissues after systemic resolution of the virus thus confirming findings in human population (15). Male to female sexual transmission of ZIKV has been observed in an interferon deficient (AG129) mouse model of ZIKV infection, resulting in a high degree of infection of AG129 females, including vertical transfer. In addition, vaginal ZIKV inoculation resulted in a high degree of infection in both non-pregnant and pregnant AG129 females. Yockey et al (16) similarly found that ZIKV replicated in the vaginal mucosa of wild type mice and while not leading to viremia, did result in fetal growth restriction and infection of the fetal brains indicating that vaginal inoculation may lead to direct transfer of the virus to the intrauterine compartment through a compromised cervical canal or via local lymphatics to the utero-placental interface. Vaginal inoculation of non-pregnant female Rhesus monkeys with ZIKV has also been achieved, albeit using raw virus in culture media diluent rather than semen in the inoculant. A recent study suggested that human semen may actually inhibit ZIKV infectivity in an *in vitro* setting, including human reproductive tract explants (17). Studies estimate sexual transmission of ZIKV in human population to be responsible for substantial number of infections and maintenance of virus in human population with or without the presence of the vector. This is particularly of concern in pregnant women as studies have shown ZIKV to be disseminated to placenta and fetus after intravaginal infections and cause CZS in fetus (7, 16, 18–20). Sexual transmission is also a likely means for the global spread of the virus between land masses.

The present study focused on the time course of the emergence and persistence of ZIKV in the blood, saliva, cervico-vaginal washings, and in maternal reproductive and fetal tissues following intravaginal inoculation using ZIKV containing semen in baboons at mid-gestation. In addition, we compared responses to both French Polynesian and Puerto Rican ZIKV isolates to assess for differences in virulence, tissue tropism and vertical transfer. Although the initial link between vertical transfer of ZIKV and microcephaly was first described in the French Polynesian outbreak after *post-hoc* analysis, a dramatic increase in microcephaly and CZS was observed in Brazil and other tropical American regions including Puerto Rico. Although an initial examination of isolates found a single point mutation in the FP isolate to likely contribute to neuroprogenitor cell targeting by ZIKV and potentially vertical transfer mechanisms, American isolates have accumulated further mutations and thus may render a more severe pregnancy outcome. This study targets many gaps in current knowledge about viremia and pregnancy outcome due to ZIKV infection transmitted sexually in a highly relevant non-human primate, the Olive Baboon.

## MATERIALS AND METHODS

### Ethical Statement

All experiments utilizing baboons were performed in compliance with guidelines established by the Animal Welfare Act for housing and care of laboratory animals as well as the United States National Institutes of Health Office of Laboratory Animal Welfare Public Health Service Policy on the Humane Care and Use of Laboratory Animals in Assessment and Accreditation of Laboratory Animal Care (AAALAC) International and National accredited laboratories. All experiments were conducted in accordance with and approval from the University of Oklahoma Health Sciences Center Institutional Animal Care and Use Committee (IACUC; protocol no. 101523-16-039-I). All studies with ZIKV infection were performed in Assessment and Accreditation of Laboratory Animal Care (AAALAC) International accredited ABSL2 containment facilities at the OUHSC. Baboons were fed standard monkey chow twice daily as well as receiving daily food supplements (fruits). Appropriate measures were utilized to reduce potential distress, pain and discomfort. Ketamine (10 mg/kg) was used to sedate baboons during all procedures, which were performed by trained personnel. All animals received environmental enrichment. ZIKV infected animals were caged separately but within visual and auditory contact of other baboons to promote social behavior and alleviate stress. At the designated times post inoculation, the animals were euthanized according to the recommendations of the American Veterinary Medical Association (2013 panel on Euthanasia).

### Animals

Adult timed-pregnant female olive baboons (n=6) were utilized for this study. All females were multiparous with history of successful prior pregnancies. All dams used in this study were determined to be seronegative for West Nile Virus (21).

### Virus stocks, infection and sample collection

Animals were anesthetized with an intramuscular dose of Ketamine (10 mg/kg) before all procedures (viral inoculation, blood, salivary swabs and cervico-vaginal washing collection). Timed-pregnant female baboons were infected vaginally with a dose of 10^6^ focus forming units (ffu; 1 ml volume per dose) of the baboon semen-spiked with the French Polynesian ZIKV isolate (H/PF/2013) or the Puerto Rican ZIKV isolate (PRVABC59) placed at the cervical os using a speculum to aid with deposition. Semen samples were collected by rectal probe ejaculation as previously described for our laboratory (15) from total of 11 male baboons seronegative for WNV and ZIKV and stored immediately at -80°C. Each semen aliquot was thawed on ice before adding virus to it for vaginal infection. Semen samples were chosen at random per inoculation. Inoculations were repeated every 7 days until ZIKV RNA was detected in whole blood by qPCR, or in one dam that never became viremic, through 4 weekly inoculations. The dosage used to infect the animals in our study is based on the previous works done in mosquitoes carrying WNV and DENV, where it was estimated that mosquitoes carry 1×10^4^ to 1×10^6^ plaque forming units (PFU) of the virus (22), from a study evaluating Brazilian ZIKV in a bite from *Aedes aegypti* mosquito and from a study of mosquito transmission of ZIKV in rhesus monkeys (23). The pregnant females were infected near mid-gestation (between 86 and 95 days of gestation [dG]; term is approx. 181 dG). Maternal blood and saliva samples were obtained on the day of inoculation (day 0) as well as at 4, 7, 11, 14, 21, and 28 days post infection (dpi). Cervico-vaginal washings were completed utilizing 3 mL of normal saline loaded into a 5 mL syringe with a catheter tip then injected onto the cervix and posterior vaginal fornix and subsequently recollected. These were obtained on days -4 (pre-wash), 4, 11, 14, 21, and 28 post infection. Ultrasound evaluation of fetal viability was completed with each inoculation and inter-inoculation specimen collection. Whole blood was collected into EDTA tubes. Saliva and vaginal samples were collected by cotton roll salivette. The sampling procedure for each dam is detailed in **Table 1**.

**Table 1.**
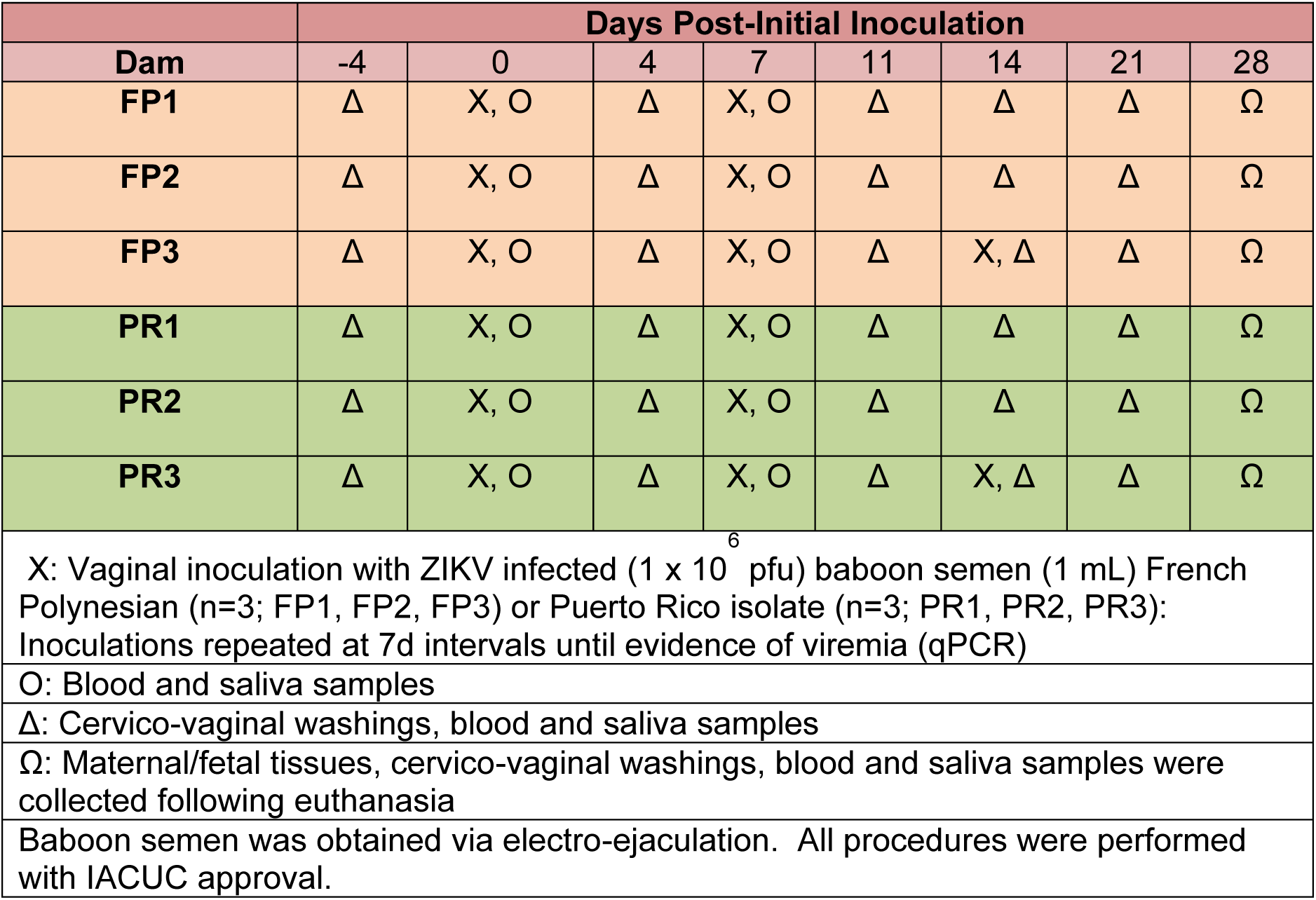
Inoculation and Sampling Procedures

At the end of the study for each animal, dams were sedated with ketamine, all maternal samples obtained as well as ultrasound measurements, then the animal rapidly euthanized with euthasol. A C-section was quickly performed, cord blood obtained and the fetus euthanized with euthasol. Maternal and fetal tissues were rapidly collected and samples were both fixed with 4 % paraformaldehyde and frozen on dry ice (stored at −80°C) for each tissue.

### Complete blood counts (CBCs)

CBCs were obtained for all females on EDTA-anticoagulated whole blood samples collected on day 0 and subsequent days-post infection as shown in the experimental timeline (Idexx ProCyte DX hematology analyzer; Idexx laboratories, ME). CBCs included analysis for red blood cells (RBCs), hemoglobin, hematocrit and platelet count.

### One-step quantitative reverse transcription PCR

Primers and probes used for qRT-PCR were designed by Lanciotti et al (24) (**Table 2**). RNA was isolated from maternal and fetal tissues (**Table 3 and 4**) using QIAamp cador pathogen mini kit (Qiagen, Valencia, CA). ZIKV RNA was quantitated by one-step quantitative real time reverse transcription PCR using QuantiTect probe RT-PCR kit (Qiagen) on an iCycler instrument (BioRad). Primers and probes were used at a concentration of 0.4 μM and 0.2 μM respectively and cycling conditions used were 50°C for 30 min, 95°C for 15 min followed by 40 cycles of 94°C for 15 s and 60°C for 1 min.

**Table 2.**
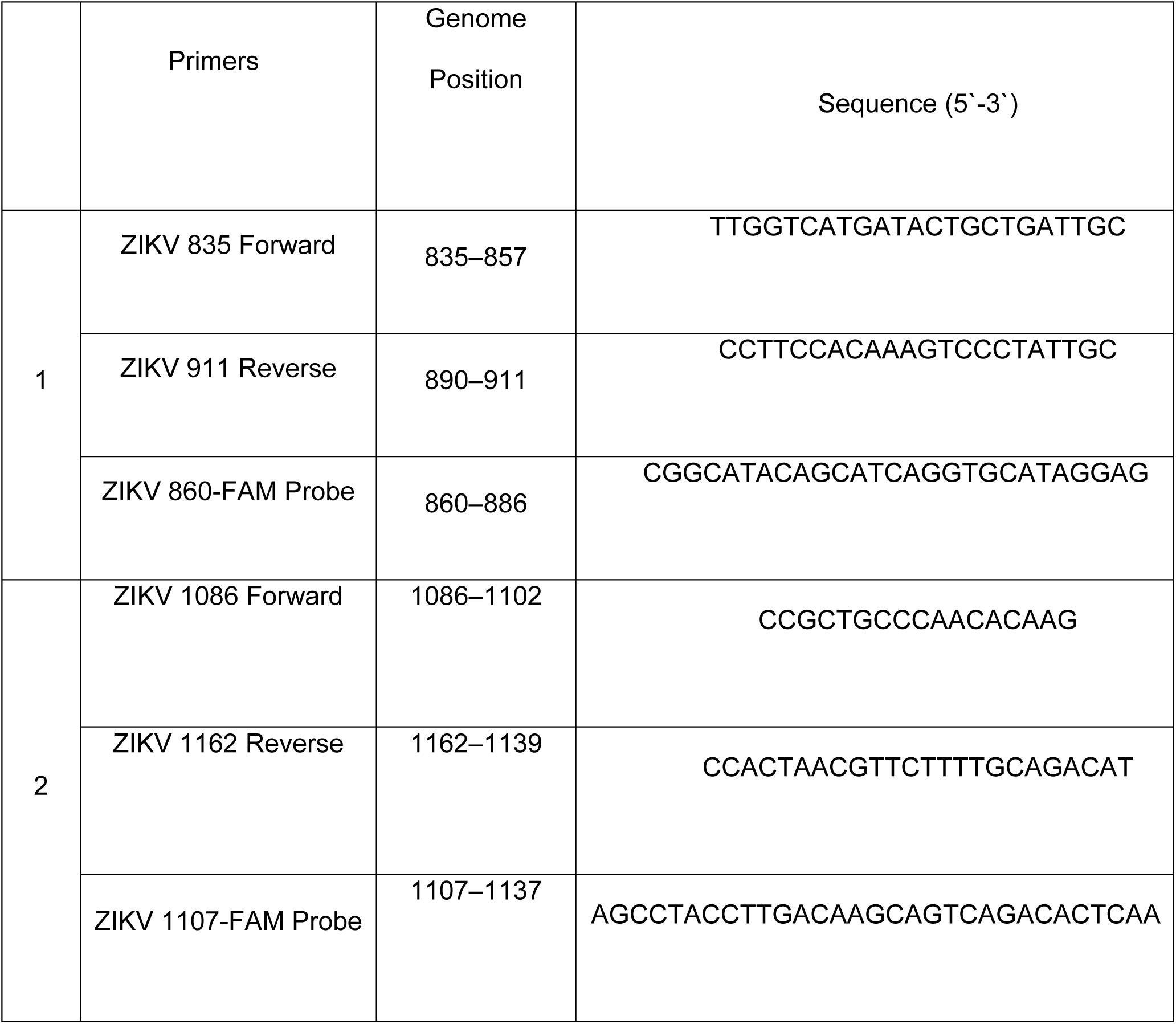
Primer/Probe sets for the detection of ZIKV by one step qRT-PCR.

**Table 3.** ZIKV RNA (log) – Maternal Reproductive Tissues and Maternal Lymph nodes.

|  | Maternal Reproductive Tissues – ZIKV RNA (log) |  |  |  | Maternal Lymph Nodes – ZIKV RNA (log) |  |  |
| --- | --- | --- | --- | --- | --- | --- | --- |
| FRENCH POLYNESIAN (FP) ISOLATE |  |  |  |  |  |  |  |
| Dam | Uterus | Cervix | Vagina | Ovary | Inguinal | Mesenteric | Axial |
| FP1 | - | - | - | - | 2.5E05 | 3.3E06 | 1.0E05 |
| FP2 | - | - | - | - | 7.6E05 | 2.0E05 | 5.4E05 |
| FP3 | - | - | - | - | 7.7E03 | - | 5.0E03 |
| PUERTO RICAN (PR) ISOLATE |  |  |  |  |  |  |  |
| PR1 | 3.2E03 | - | 6.9E03 | - | 2.7E03 | - | 4.1E03 |
| PR2 | 4.7E04 | - | - | - | 2.7E05 | 5.8E05 | 3.2E05 |
| PR3 | - | - | 2.8E03 | - | 1.2E04 | 7.1E04 | - |
| - : below level of detection |  |  |  |  |  |  |  |

Concentration of the viral RNA (copies/milliliter) was determined by interpolation onto a standard curve of six 10-fold serial dilutions (10^6^ to 10^1^ copies/ml)) of a synthetic ZIKV RNA fragment available commercially from ATCC (ATCC VR-3252SD). The cutoff for limit of detection of ZIKV RNA was 1×10^2^.

## ZIKV ELISA

ZIKA specific IgM and IgG antibody responses were assessed in the serum samples using the commercially available anti-ZIKV IgM (#ab213327, Abcam, Cambridge, MA) and IgG (#Sp856C, XpressBio, Fredrick, MD) ELISA kits. Briefly, a 1:100 for IgM and 1:50 for IgG serum dilution was performed in duplicate and added to the pre-coated plates available in the kits. The assays were performed using the manufacturer’s instructions and the assay was read at 450 nm for IgM and 405 nm for IgG antibodies in the serum.

### Immunofluorescence

For IF, slides were baked for one hour at 56°C, deparaffinized, and HIER was performed in the Retriever 2100 with R-Universal Epitope Recovery Buffer (62719-10 lot 180314. After retrieval, slides were blocked in 5% normal donkey serum for 1 hour, then primary antibodies in 0.5% normal serum were added and incubated overnight, humidified, at 4°C [MAC-387, macrophage antibody (Abcam, MA); Pan anti-flavivirus; (Millipore, CA)]. The next morning slides were removed from 4°C and allowed to equilibrate to RT, covered, on the benchtop for 1 hour. Slides were rinsed 4 x 5 minutes with PBS, then secondary antibodies were added and incubated 1 hour, covered, at RT. Donkey anti-mouse IgG F(ab’)2 AlexaFluor 594 (Jackson Immunolabs) was used as secondary antibody. Slides were rinsed in PBS, counterstained 5 minutes with DAPI in PBS and cover slipped using Shur/Mount. Cover glass were sealed with nail polish and slides were stored at 4°C and visualized using a fluorescent microscope (Olympus BX40). Images were captured using CellSens imaging software (Olympus).

## RESULTS

### Description of animal cohorts and experimental outline

For this study, adult timed pregnant female olive baboons (n=6) were used. All baboons were infected via vaginal inoculation with a clinically relevant dose (1×10 ^6^ pfu) of the French Polynesian (H/PF/2013) or the Puerto Rican (PRVABC59) ZIKV isolate. Blood, saliva, and cervico-vaginal washings were collected as shown in **Table 1**. In the FP cohort, 2/3 dams developed slight to negligible rash on the abdomen and in the inguinal and axillary regions and no conjunctivitis. One FP dam (FP1) presented with moderate rash on the abdomen, in the bilateral axillary and inguinal regions and mild conjunctivitis. In the PR cohort, 2/3 dams developed slight to negligible rash on the abdomen and in the axillary/inguinal regions and only one dam (PR3) developed slight conjunctivitis. None of the animals showed signs of any other clinical disease. Baboons were euthanized at 28 dpi. Complete blood counts (CBCs) were evaluated for all females on EDTA-anticoagulated whole blood samples collected on day 0 and subsequent days post infection as shown in Table 1 (Idexx ProCyte DX hematology analyzer; Idexx laboratories, ME). RBC, hemoglobin and hematocrit numbers did not show any differences pre-and post ZIKV infection in all females (data not shown). Platelet counts did not change in response to ZIKV infection in any dam (data not shown).

### Viral load data post infection in whole blood and saliva

Viral RNA was quantified by one-step qRT-PCR in RNA extracted from the blood and saliva samples. ZIKV RNA was detected in the blood of all animals infected with the FP isolate of ZIKV and two of the three animals infected with the PR isolate of ZIKV (**Figure 1**). Among the dams infected with the FP ZIKV isolate, ZIKV RNA was detected in the blood of all three dams, with dam FP1 viremic at 7 and 11 dpi, dam FP2 viremic at 4 and 7 dpi and dam FP3 viremic at 4 and 7 dpi (**Figure 1A**). Among the dams infected with the PR ZIKV isolate, ZIKV RNA was detected in the blood of 2 of 3 dams; dam PR1 was viremic at 7, 11, 14 and 28 dpi, dam PR2 was viremic on 4, 7 and 14 dpi. ZIKV RNA was never detected in the blood of dam PR3 at any time point examined (4, 7, 11, 14, 21, 28 or 35 dpi) (**Figure 1B**).

**Figure 1.**
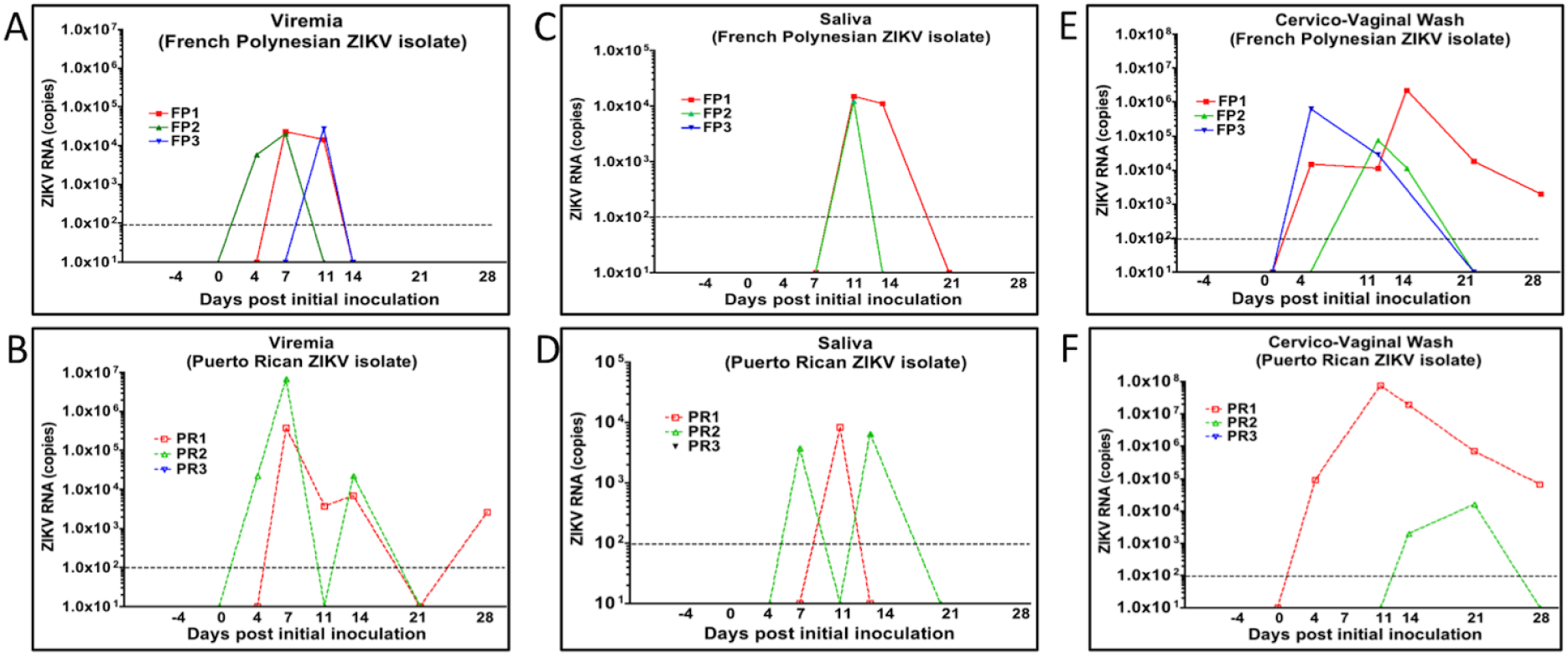
ZIKV RNA in blood (viremia, A, B), saliva (C, D) or cervico-vaginal washes (E, F) in mid-trimester baboons inoculated intra-vaginally with baboon semen containing either the FP or PR ZIKV isolates. ZIKV RNA was prepared from specimens collected from each animal at the indicated days post-infection and quantitated by one-step qRT-PCR.

In the two dams infected with the FP ZIKV isolate, ZIKV RNA was detected in the saliva at 11 and 14 dpi (FP1) and on 11 dpi (FP2) (**Figure 1C**). In the two animals infected with the PR ZIKV isolate, ZIKV RNA was detected in the saliva at 11 dpi (PR1) and 7 and 14 dpi (PR2) (**Figure 1D**).

### ZIKV shedding into cervico-vaginal washes (CVW)

ZIKV RNA was detected in the CVW of all three dams infected with the FP isolate of ZIKV and two of the three dams infected with the PR isolate of ZIKV. ZIKV RNA was detected in the CVW at 4, 11, 14, 21 and 28 dpi in dam FP1, 11 and 14 dpi in dam FP2 and 4 and 14 dpi in dam FP3 (**Figure 1E**). In the PR ZIKV infected dams, two had ZIKV RNA in the CVW; 4, 11, 14, 21 and 28 dpi for PR1 and 14 and 21 dpi for dam PR2 (**Figure 1F**).

### ZIKV RNA in maternal tissues

In maternal reproductive tissues (uterus, cervix, vagina, ovaries), ZIKV RNA was not detected in any of the animals inoculated with the FP ZIKV isolate. ZIKV RNA was detected in the uterus of two of the animals inoculated with the PR isolate (dams PR1, 2) and in the vagina of two dams (PR1, 3). ZIKV RNA was present in all of the maternal lymph nodes assessed (axial, mesenteric, inguinal) except for the mesenteric nodes of dam FP3 and the axial lymph nodes of dam PR3 (**Table 3**).

### ZIKV shedding into fetal tissues and placenta

ZIKV RNA was not detected in any of the fetal tissues (cord blood, cortex, cerebellum, umbilical cord, fetal membranes, spleen, lung, liver, eye, gonads, stomach, intestine, and optic nerve; data not shown).

Placentas from each dam were sampled from six different locations (different cotyledons). In the animals infected with the FP ZIKV isolate, ZIKV RNA was not detected in any cotyledons sampled. In the animals infected with the PR isolate, ZIKV RNA was detected in two of the animals (PR1, 2). In one of these animals, ZIKV RNA was detected in five cotyledons sampled (PR1), and in the other animal, ZIKV RNA was detected in four cotyledons sampled (PR2) (**Table 4**).

**Table 4.**
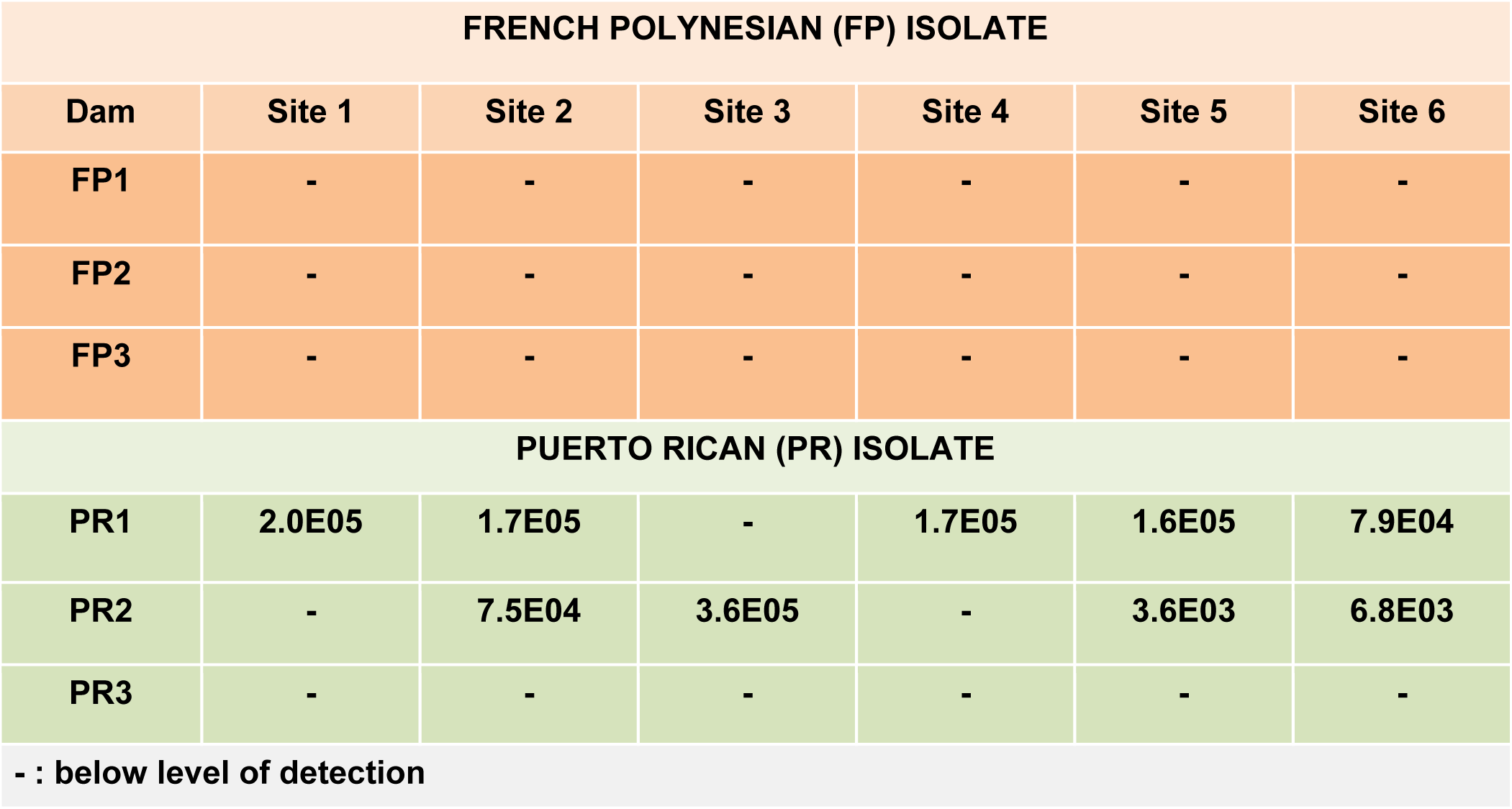
ZIKV RNA (log) – Placental Tissues.

### ZIKV antibody response

ZIKV IgM was detected in sera of 2 of 3 FP ZIKV isolate infected dams (FP1, 2) on day 14 and in sera of dam FP3 by day 21 post infection (**Figure 2A**). ZIKV IgM was detected in 2 of 3 PR isolate dams (PR1, 2) by day 11 post infection and in dam PR3 on day 18 post infection (**Figure 2B**). ZIKV IgG was detected in sera of dams FP 1 and 2 on day 21 and by day 35 post infection in the sera of dam FP3 (**Figure 2C**). In the PR isolate cohort, dams PR1 and 2 had detectable ZIKV IgG in sera on day 21 post infection and day 32 post infection in dam PR 3 (**Figure 2D**).

**Figure 2.**
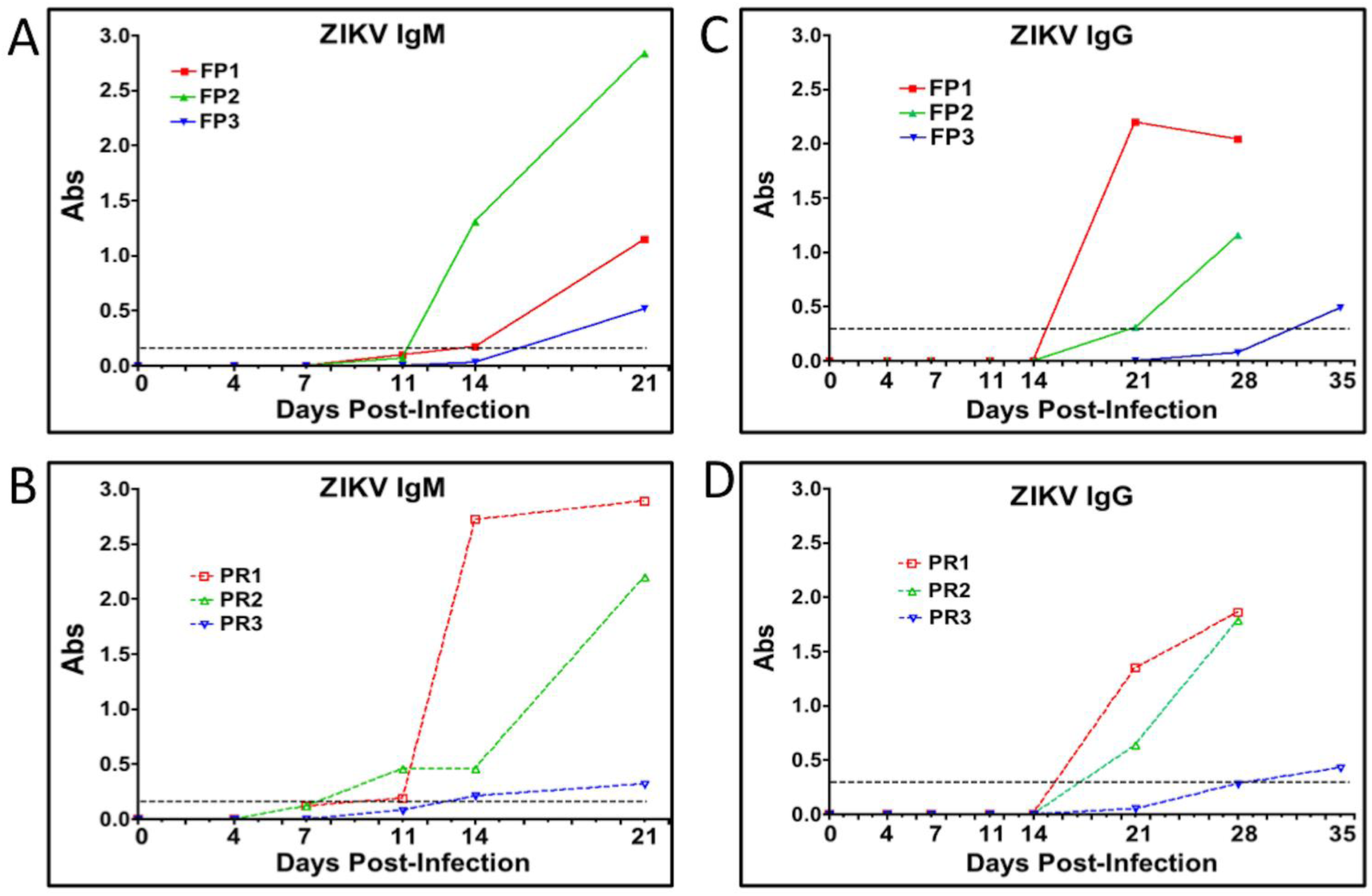
Serum ZIKV IgM (A, B) and IgG (C, D) at the indicated days post-infection in mid-trimester baboons inoculated intra-vaginal with baboon semen containing either the FP or PR ZIKV isolates. The dashed lines represent the assay cutoff controls for IgM and IgG detection in each specific ELISA.

### Immunohistochemistry

#### Cervix

ZIKV immunofluorescence (IF; pan flavivirus) was observed in the cervix of one dam infected with the French Polynesian ZIKV isolate (FP1) but not in dams FP2 or FP3. In FP1, the ZIKV IF pattern was localized to the epithelial layer in both the endocervical (upper cervix) and ectocervical (lower cervix, exterior os) regions. In the simple columnar epithelium characteristic of the endocervix, the strongest IF resided near the basil lamina separating the epithelium and stromal layers with diffuse IF observed in the epithelial layer (**Figure 3A**). In the ectocervix, the stratified squamous epithelium exhibited ZIKV IF in cells throughout the epithelium from the surface to the basal lamina cell layer. No ZIKV IF was noted in this dam in the cervical stroma. ZIKV IF was observed in the cervix of 3/3 dams infected with the Puerto Rican ZIKV isolate (**Figure 3**). In these dams, ZIKV IF was observed in the epithelial layer of both the endo- and ectocervix with a more intense IF compared to the dam infected with the French Polynesian isolate. In addition, occasional cells exhibited ZIKV IF in the cervical stroma, potentially representing either isolated stromal cells or invading immune cells such as macrophages or neutrophils. In addition, an occasional cervical gland was observed to exhibit ZIKV IF (**Figure 3**).

**Figure 3.**
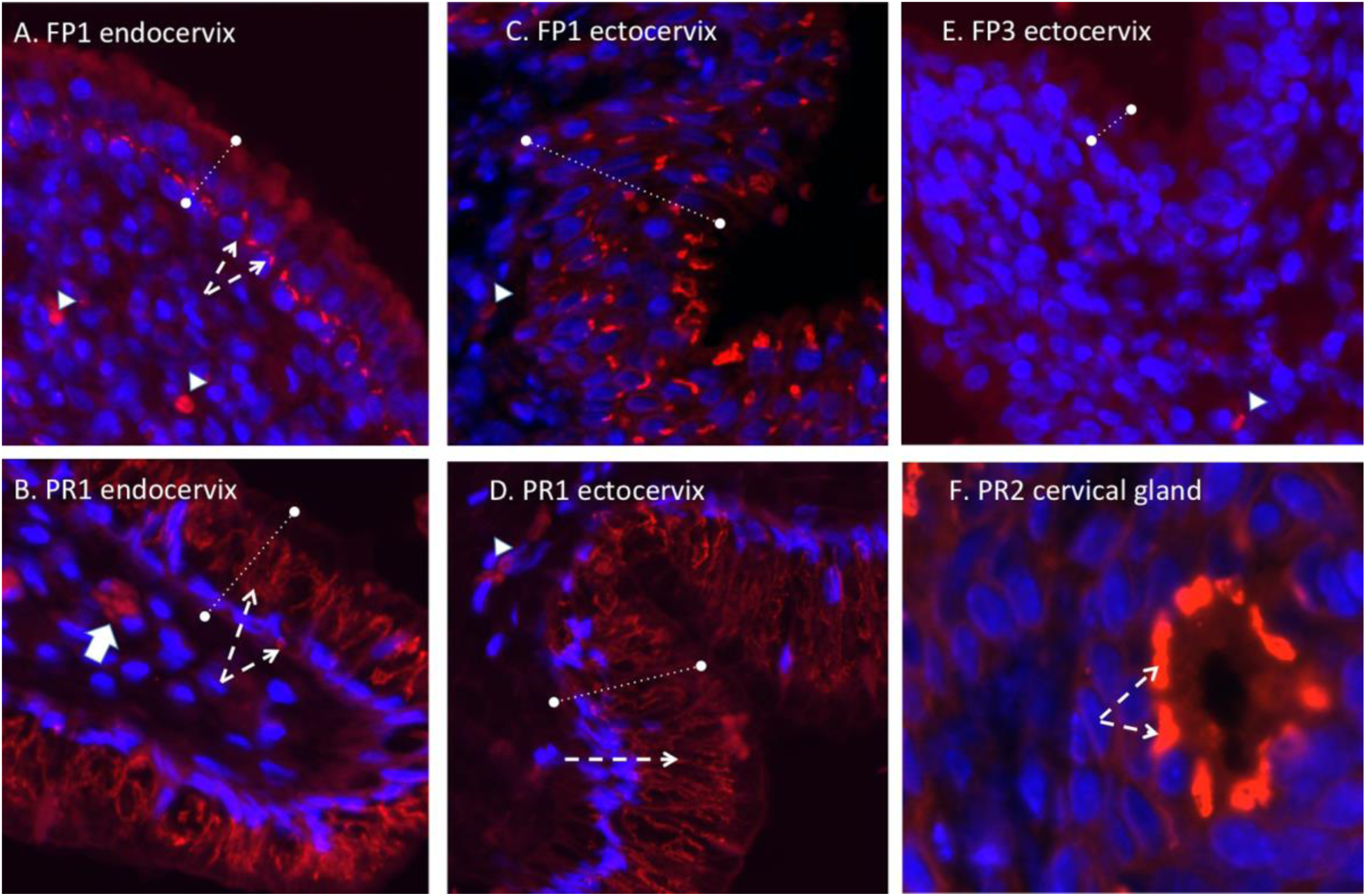
Pan flavivirus immunofluorescence (IF; panflavi, Red; DAPI, Blue) in the cervix. In the endocervix (A, B; upper cervix) ZIKV IF (dashed white arrows) was localized in the epithelium (denoted by the dashed lines) in both the endocervical region (A,B; simple columnar epithelium) and the ectocervix (CD; lower cervical-vaginal region, stratified squamous epithelium) in both PR and FP inoculated dams. In the FP isolate inoculated dam (FP1), intense ZIKV IF was observed at the interface of the epithelial-stromal junction in the endocervical region (A, B) with a broader distribution in the epithelium in the ectocervical region (C, D). In the PR isolate infected dams (B, D) a more intense ZIKV IF staining was noted in the epithelium of both the cervical regions as well as in occasional cells in the stroma (white arrow, B). In one PR isolate infected dam, ZIKV IF was also noted in the cervical glands (F, dashed white arrows). For comparison, a dam infected with the FP isolate (E) that did not exhibit ZIKV IF in either cervical region is shown. Small arrowheads denote auto-fluorescing red blood cells.

In the cervix of the French Polynesian and Puerto Rican ZIKV isolate infected dams that exhibited cervical ZIKV IF, macrophages were frequently observed in the stromal layer juxtaposed to the epithelial layer (**Figure 4**). Puerto Rican isolate infected dams (PR1, 2) exhibited a greater infiltration of macrophages into the stroma compared to FP1 with macrophages being observed both adjacent to the epithelial layer as well as deeper in the stromal layer (**Figure 4C**). In the French Polynesian isolate infected dams that did not exhibit cervical ZIKV IF, only occasional macrophages were observed in the stroma (**Figure 4D**).

**Figure 4.**
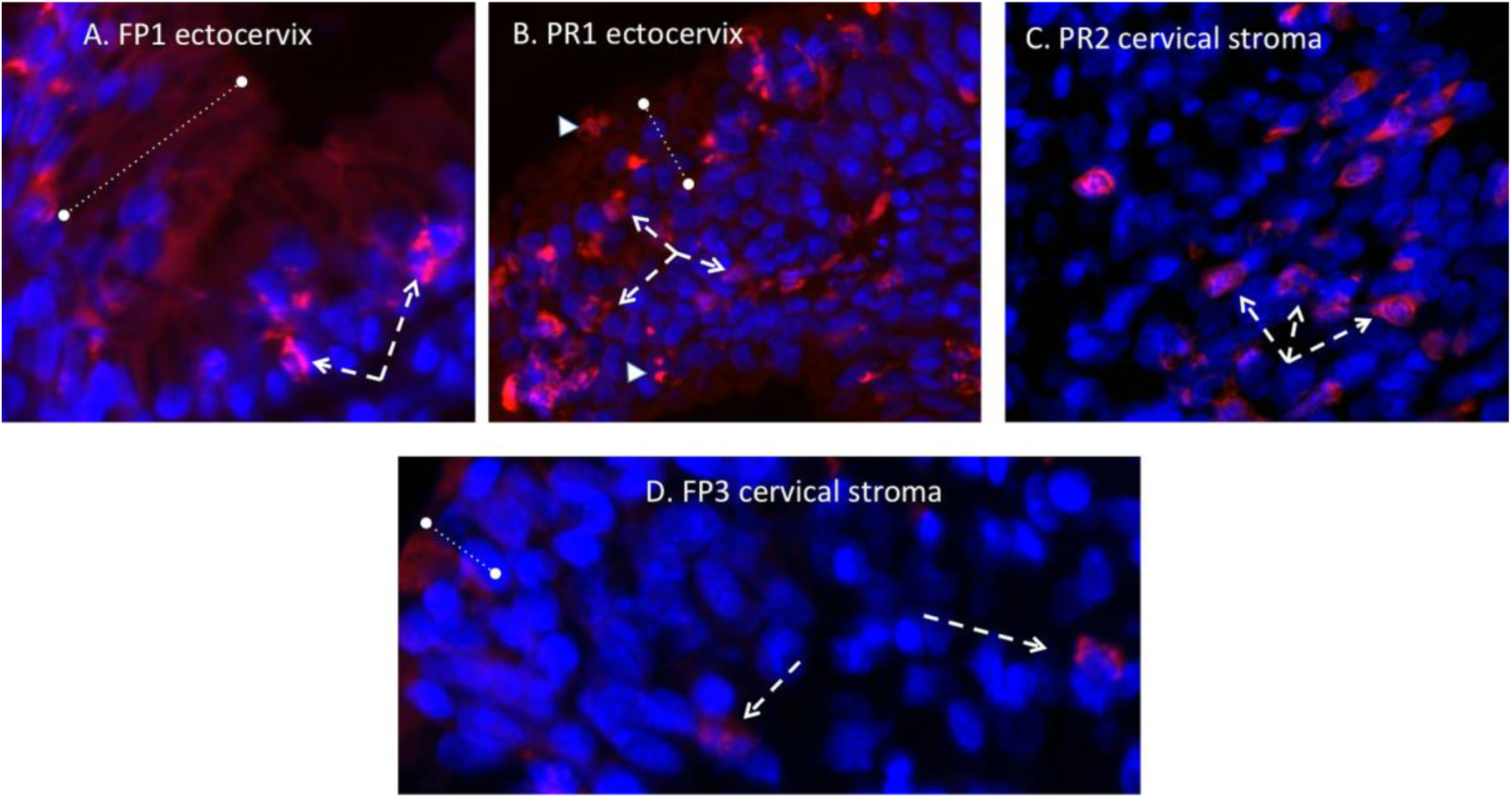
Immunofluorescent staining in the cervix for macrophages (Red; DAPI: Blue). In both FP isolate and PR isolate infected dams, abundant macrophages were observed, in the FP isolate infected dam that exhibited ZIKV IF (A, FP1), macrophages (dashed arrows) were primarily localized at or near the epithelial-stromal interface. In the PR isolate dams that were ZIKV IF positive, macrophages were localized at the both the epithelial-stromal interface region (B) as well as deeper in the stomal tissue (C). In dams that exhibited no ZIKV IF in the cervix, (D), only occasional macrophages were noted, typically in the deeper stomal layer.

#### Uterus

No Zika virus (pan flavivirus) IF was observed in the myometrium of any dam infected with either French Polynesian or Puerto Rican ZIKV isolate (**Figure 7**). Occasional clusters of ZIKV IF positive cells were observed in the endometrium of the two dams infected with the Puerto Rican isolate that were ZIKV RNA positive in the uterus, but not in the dams infected with the French Polynesian isolate or the one dam infected with the Puerto Rican isolate that did not have detectable ZIKV RNA in the uterus. We did not detect macrophages in any of the uterine sections examined from any animal.

**Figure 7.**
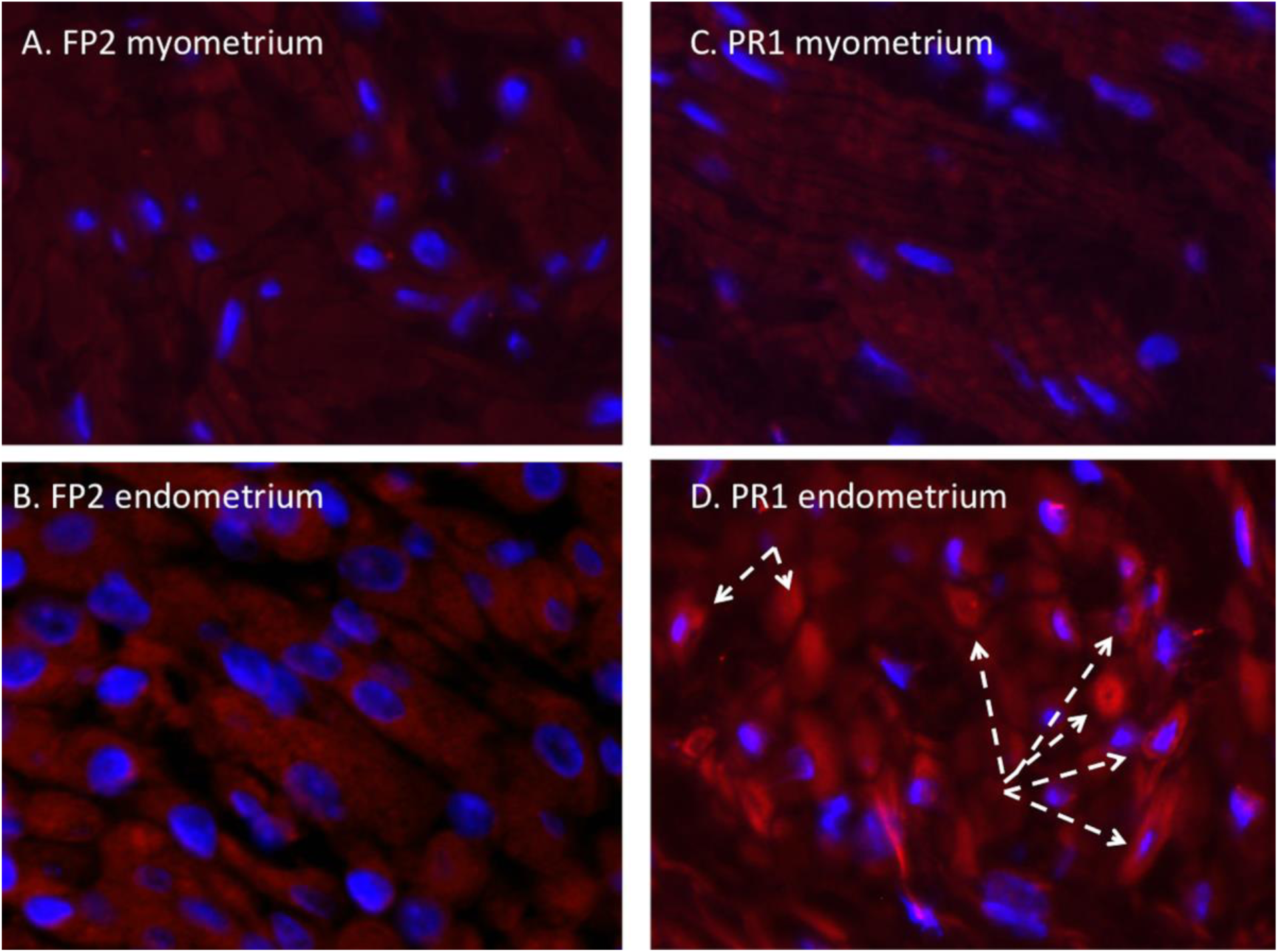
Immunofluorescent staining in the uterus for ZIKV (Red; DAPI: Blue). ZIKV IF was not observed in the uterus, either endometrium or myometrium of FP isolate infected dams (A, B). In the PR isolate infected dams, ZIKV IF was not observed in the myometrium and was occasionally observed in clusters of endometrial cells, (dashed arrows, D).

#### Placenta and fetal membranes

The dams infected with the French Polynesian ZIKV isolate were negative for ZIKV IF in the placenta and fetal membranes. ZIKV IF (pan flavivirus) in the placentas of Dams PR1 and PR2 (both positive for placental ZIKV RNA) demonstrated the presence of ZIKV, localized primarily in the syncytial layer with regions exhibiting greater intensity (**Figure 5**). In these two dams, we observed ZIKV IF in the amnion epithelium (PR1) and in both the amnion and chorion/decidua of the fetal membranes. Of note, the fetal membranes examined in this study were adjacent to the utero-placental interface and not the more distal membranes. We did not observe macrophage IF in these sections (**Figure 6**).

**Figure 5.**
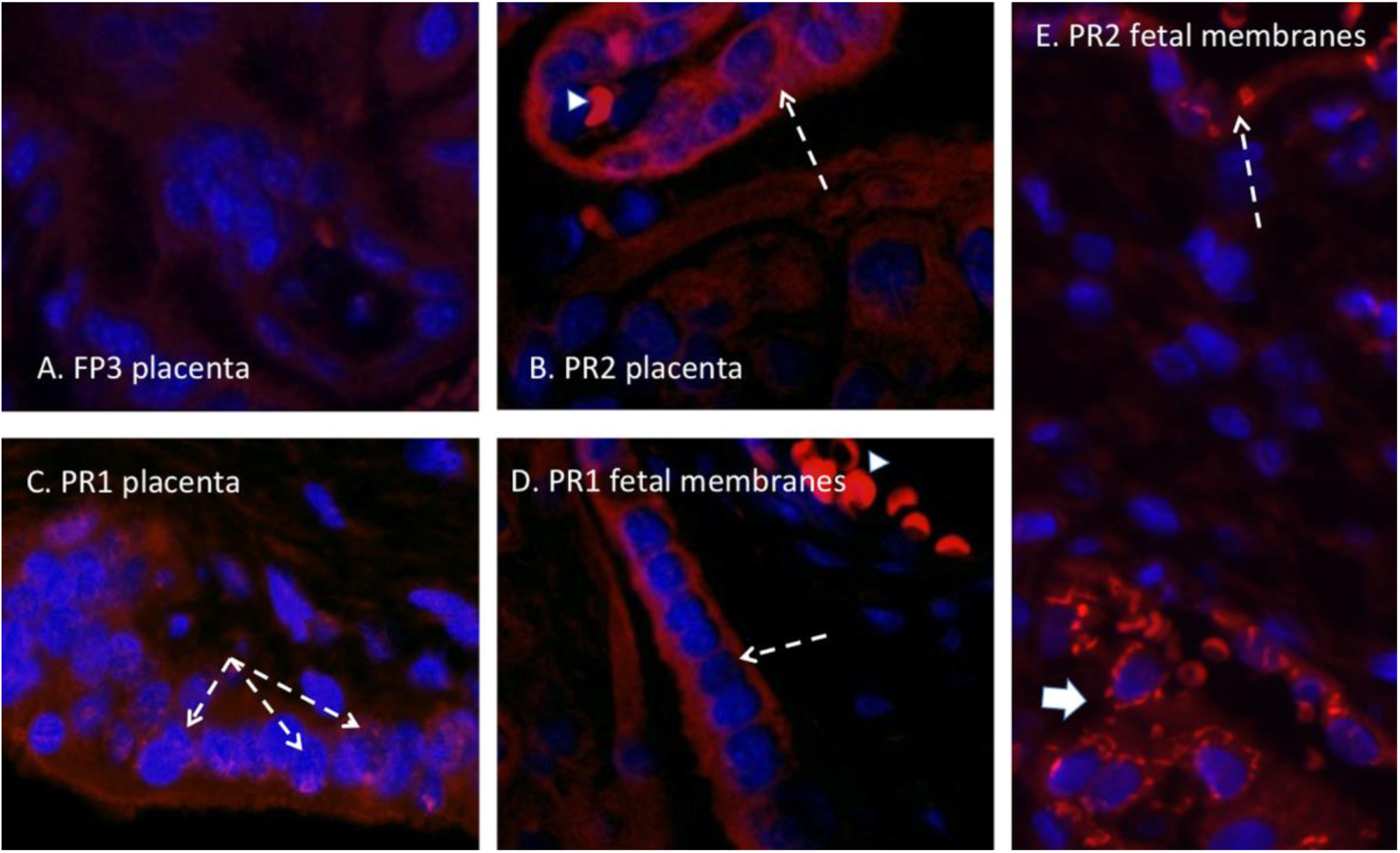
Pan flavivirus immunofluorescence (IF, Red: flavivirus; blue: DAPI) staining in the placenta from a ZIKV RNA negative placenta from a dam infected with the FP isolate (A) demonstrating a lack of ZIKV IF in villous placenta. ZIKV IF was noted in occasional villous trophoblast in placenta from dams infected with the PR isolate (B, C; white arrows) consistent with ZIKV infection of the syncytial layer. Infection of the amnion epithelium was also observed (D; arrow). ZIKV IF was also observed in the chorion/decidual layer of the fetal membranes in a dam infected with the PR isolate (E. large arrow). Auto-fluorescing red blood cells are indicated with arrow heads.

**Figure 6.**
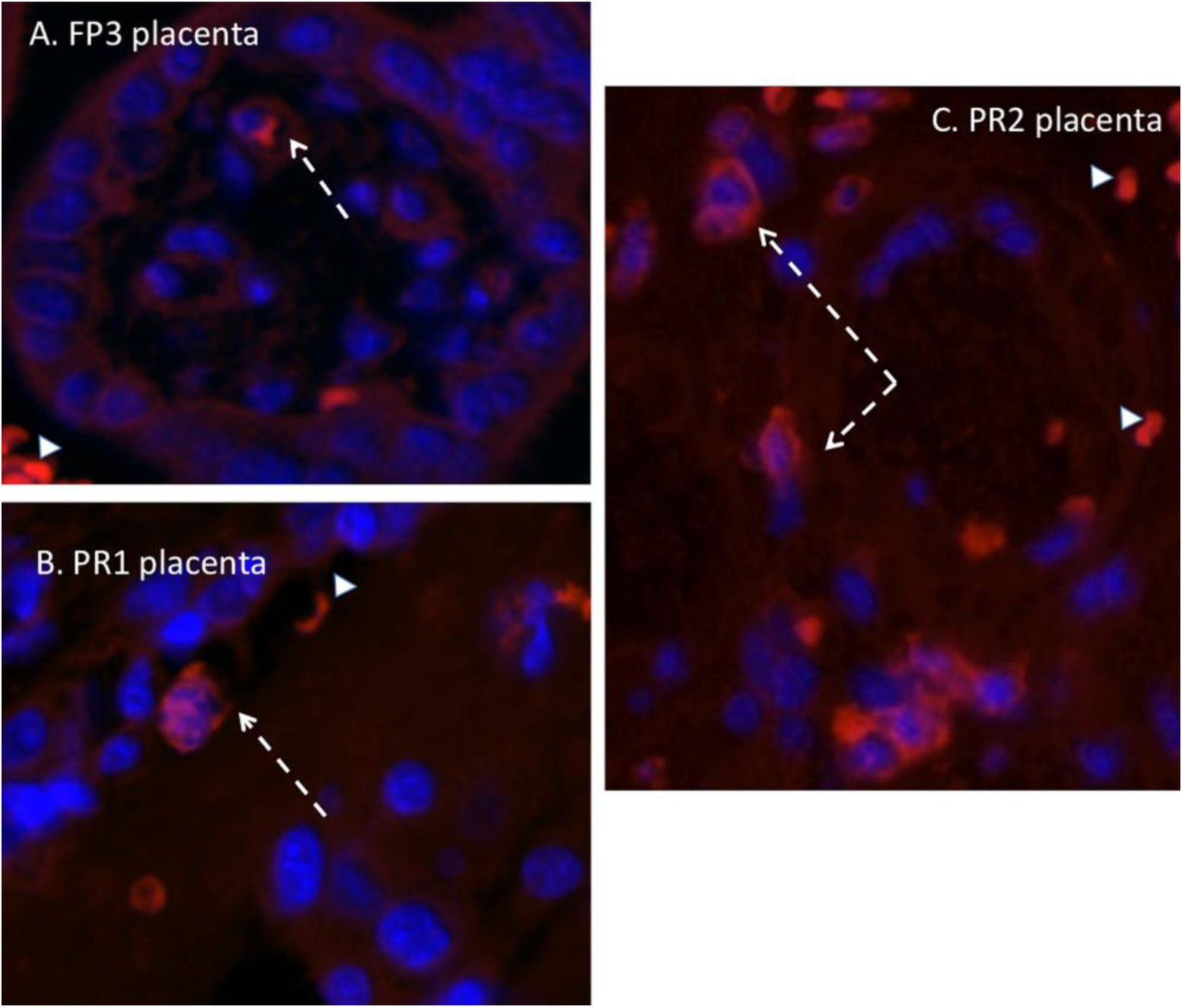
Immunofluorescent staining in the placenta for macrophages (Red; DAPI: Blue). In both FP isolate infected dams, only an occasional macrophage was observed in the maternal and fetal compartments of the placenta (A: arrow). In the PR isolate dams that were ZIKV IF and RNA positive, macrophages were more abundant, in particular in the maternal compartment of the placenta (C: arrow). Arrowheads denote auto-fluorescing red blood cells.

## DISCUSSION

In the present study, we describe the ZIKV infection of six timed-pregnant olive baboons at mid-gestation (86-95 dG; term ∼183 dG) following vaginal deposition of 1 mL baboon semen containing 1 x 10^6^ pfu of either the FP or the PR isolates of ZIKV. We chose the FP isolate based on our prior study where we demonstrated vertical transfer following subcutaneous inoculation with the FP isolate that resulted in both fetal demise as well pronounced fetal CNS pathology (25). A retrospective study of the ZIKV epidemic in French Polynesia (circa 2013) noted that this was the first instance associating ZIKV to microcephaly and congenital ZIKA syndrome (CZS) (21, 26). It was later reported that the FP isolate differs from the ancestral Asian ZIKV lineage from which it was derived, with a mutation in the prM protein (S139N), which has been stably maintained throughout the virus’s dissemination throughout the Americas (Fig 8). This mutation is associated with enhanced infectivity in human neural progenitor cells (NPCs) and yielded a more significant microcephaly in mice (27). We selected the PR isolate to compare to the FP isolate to examine for increased rates of vertical transfer, since it harbors the S139N mutation and has also acquired several additional point mutations resulting in amino acid substitutions, some being common with the Brazilian isolate(s) (28). Similar to the increased incidence of CZS noted in the Brazilian ZIKV epidemic (compared to the French Polynesian estimates), a recent study by CDC reported that one in seven children born from women with confirmed or possible ZIKV infection during gestation in Puerto Rico had a birth defect or neurodevelopmental abnormality suggesting possible mutations in this ZIKV isolate may contribute to the increased viral replication and neurovirulence compared to the FP strain (29).

**Figure 8.**
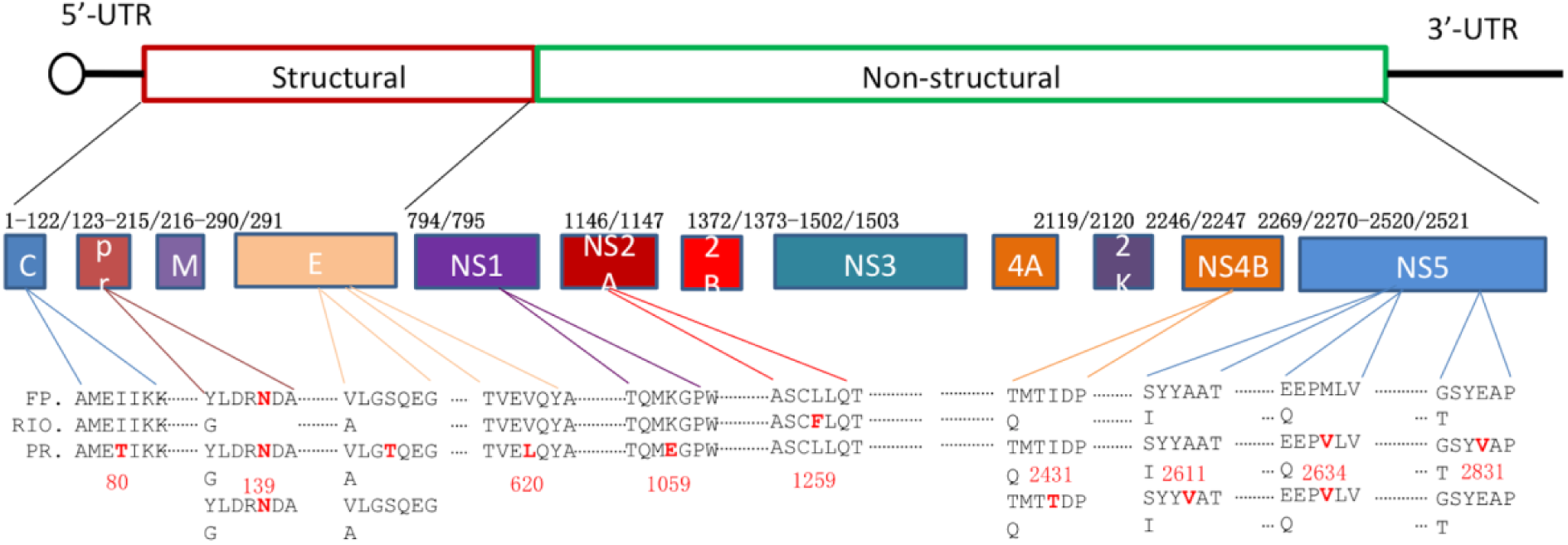
Amino acid variance among Zika virus isolates. The amino acid sequences for the FP (H/PF/2013; AHZ13508), PR (PRVABC59; AYI50388), and RIO (RIO-U1/2016; AMD16557) complete polyproteins were aligned using the Clustal Omega algorithm in Geneious Prime 2020.0.3 (www.geneious.com). Graphical representation of the ZIKV genome as well as the resulting protein products is indicated. Amino acid variances are highlighted in red with their position in the polyprotein noted by numerical annotation as well as the nonstructural protein where they reside. The S139N mutation in the prM protein is noted in all isolates for reference.

In the current study, ZIKV inoculations were repeated at seven-day intervals until viremia was evident via qPCR or through three inoculations if no viremia was observed (one PR infected dam). We chose multiple inoculations to mimic probably repeat intercourse in human couples. Viremia was achieved in all dams except one of the three baboons inoculated with the PR isolate noted between days 4 and 14 post infection (Fig 1). In those animals inoculated with the FP isolate, resolution was noted by day 14 post infection, while in dams inoculated with the PR isolate, one dam had resolution by day 11 post infection with reemergence at day 14 post infection with resolution noted again at day 21 post infection (PR 2). The other PR dam (PR 1) had resolution by day 21 post infection and reemergence at day 28 post infection. The course of viremia in response to vaginal delivery of ZIKV infected semen differed from that described in our previous study in mid-gestation pregnant baboons subcutaneously inoculated with FP isolate of ZIKV where viremia was resolved by 14 dpi (25). Therefore, the route of infection with ZIKV seems to affect the length of viremia and may even affect resolution and re-emergence of the virus. We can speculate that the re-emergence of the virus could be attributed to the virus from several reservoirs where ZIKV has been found to persist, including the gastrointestinal tract, the cerebrospinal fluid, and the lymph nodes. These tissues have been shown to persistently harbor ZIKV despite a robust immune response (21, 25, 30–32). We found persistence of ZIKV RNA in both reproductive tissues as well as lymph nodes at the end of the study, serving as possible reservoirs for viral re-entry in the present study. While the lymph nodes were found to contain ZIKV RNA at the termination of the study in both FP and PR isolate inoculated dams, only the PR dams exhibited re-emergence of viremia (Table 3). This could be either strain related from an increased virulence of the PR isolate or from the reproductive tissues since only the PR isolate inoculated dams exhibited vaginal or uterine ZIKV RNA at the study termination. Unlike the paper by Müller et al (17) describing the inhibition of ZIKV infection of cells extracted from human anogenital tract and various human reproductive tissue explants by human semen treatment, we detected ZIKV infection in bodily fluids and tissues of dams infected vaginally by semen samples from ZIKA seronegative male baboons mixed with either FP or PR ZIKV isolate. Therefore, semen did not prevent ZIKV infection in baboons inoculated vaginally with ZIKV. Several studies in mice have also shown sexual transmission of ZIKV through mating of ZIKV infected male mice with ZIKV naïve female mice and *in utero* transmission in pregnant mice due to sexual transmission (20, 33, 34).

Studies have shown extended viremia in pregnant macaques following subcutaneous inoculation where viremia is characteristically prolonged for several weeks to months (35–37). It has been proposed that the prolonged or re-emergence of viremia in macaques could be from tissues harboring ZIKV including placenta (38). It is also noteworthy for the latter that only the PR isolate inoculated baboon dams exhibited ZIKV RNA in the placenta. In humans, viremia lasting up to 53 days has been reported (24), but more typically it is short in duration, lasting three to seven days (39). We cannot, however, discount the possibility that we could have seen extended viremia in the one PR dam (PR1) post 28 dpi and/or re-emergence on a different time point in any of the dams had we extended the study for a longer duration rather than termination at 28dpi. In addition to the differences in viremia in terms of duration, resolution and re-emergence between the FP and the PR ZIKV isolates, it is of potential interest that the two dams inoculated with the PR isolate had viremia levels one to two orders of magnitude compared to the FP isolate inoculated dams (Fig 1). Whether this can be attributed to a greater capacity for infectivity by the PR isolate or simply variability in a small cohort study remains to be determined.

Despite viremia being observed in five of six dams, vaginal infection with FP and PR isolate presented with slight to negligible rash on the abdomen and in the axillary and inguinal regions with little or no conjunctivitis. Only one dam inoculated with FP isolate (FP1) presented with moderate rash and mild conjunctivitis. Dams infected with PR isolate presented with slight to negligible rash on the abdomen and bilateral axillary and inguinal regions with no conjunctivitis. One PR dam (PR3) presented with slight to mild rash on the lower abdomen, axillary and inguinal regions and slight to negligible conjunctivitis. None of the dams exhibited any other signs of decreased appetite, water intake, signs of fever or arthralgia. This lack of presentation of clinical signs after vaginal inoculation of ZIKV differed from the moderate to severe clinical signs we observed in all baboons (primarily rash and conjunctivitis) that we previously infected subcutaneously with the FP isolate in prior studies of mid-gestation pregnant baboons (24), as well as non-pregnant female baboons (21), and male baboons (15, 21). The difference in presentation of clinical signs is likely due to the different routes of ZIKV inoculation (subcutaneous vs vaginal) and is of significant clinical importance since, unlike, subcutaneous infection, ZIKV infection via sexual intercourse may be more asymptomatic than usual and therefore, evade preventive measures which could preclude diagnosis, in particular in pregnancy.

In the FP cohort, ZIKV RNA was detected continuously in CVW of dam FP1 from 4 through 28 dpi while resolving by 21dpi in dams FP2, 3. In the PR infected cohort, ZIKV RNA was detected in the CVW of dam PR1 from 4dpi through 28 dpi whereas in dam PR2, ZIKV RNA was not detected until 21dpi and resolved by 28dpi (Fig 1E,F). The delayed detection of ZIKV RNA in the wash of PR3 may reflect that this dam was not infected by vaginal semen inoculation until the 3^rd^ attempt (day 14 post-initial inoculation) and also may help explain the lack of viremia observed in this dam. The longer presence of vRNA in CVW is distinct from blood and saliva and could contribute to the re-emergence of viremia in dam PR1 (14 days non viremic between last viremic time point and re-emergence) and dam PR2 (7 days non-viremic between last viremia time point and re-emergence). However, it is noteworthy that the FP dams did not show re-emergence of viremia even though one (FP1) had prolonged virus in the CVW (28 dpi). This may be possibly related to differences in infectivity of the two isolates.

In addition to the fluid compartments, we found ZIKV RNA persisting in the lymph nodes of all animals from both FP and PR isolate cohorts. In rhesus macaques, ZIKV RNA has been detected in the lymph nodes for 5 to 6 weeks post infection (30). Presence of ZIKV in the lymph nodes is one possible route of disseminating the virus from vagina to the cervix and uterus. The lymphatic system may serve as a reservoir for ZIKV persistence as well as sites of re-emergence of the virus through viral shedding from the local lymphatic tissue, possibly on an intermittent basis, by supplying virus to glandular tissue (via local lymphatic vascular beds) leading to secretion into the bodily fluids. However, at this time, we do not know if the ZIKV found in the saliva, blood, lymph nodes, reproductive tissues or cervico-vaginal washings at the termination of the study was infectious virus.

ZIKV RNA was not detected in any reproductive tissues of the dams infected with the FP isolate. However, cervical tissue of dam FP1 was found positive for ZIKV IF staining. The CVW samples of the other two FP dams were ZIKV positive only on days 4, 11 and 14 and were not ZIKV positive by qPCR or IF staining. The CVW from dam FP1 was positive for ZIKV RNA from day 4 through 28 post infection. It is possible that our inability to detect ZIKV RNA in the cervix in this dam (and the others) is related to its restriction to the epithelial layer and not stroma. In the cervix, the majority of the tissue is stromal and our tissue sampling for RNA analysis excluded epithelium. Alternatively, since the epithelium represents such a small contribution to the total RNA from cervix (vs stroma) that ZIKV RNA was below the exclusion limit for detection (1×10^2^ copies) that we set. In the PR isolate cohort, the cervix of all three dams were positive for ZIKV IF. Dams PR1 and 3 had detectable ZIKV RNA in the vaginal tissue. It is noteworthy that the vagina, which is continuous with the cervix, differs from the cervix in that it has a large stratified squamous epithelium that may help in harboring the virus. Dams PR1 and 2 were positive for ZIKV (RNA and protein) in the uterus as well, and positive ZIKV IF staining in the fetal membranes and placenta of the same two PR infected dams suggests possible spread of ZIKV to the fetus per se had the study been for a longer duration. Dams PR1 and 2 had prolonged detection of ZIKV RNA in CVW samples possibly explaining positive ZIKV IF in the cervical tissue. Although CVW samples of dam PR 3 were negative for ZIKV RNA, cervical tissue was positive for ZIKV IF staining and ZIKV RNA was detected in the vaginal tissue. It is possible that ZIKV in this animal was restricted to select tissues such as the cervix, vagina and lymph nodes and remained tissue localized rather than replicate and spread as suggested by the lack of viremia after three inoculations and weak IgM and IgG response to ZIKV. The presence of ZIKV virus in different reproductive tissues after vaginal transfer of the virus suggests that the reproductive organs in baboons may harbor ZIKV during the acute phase of ZIKV infection through sexual transmission.

Recruitment of macrophages into the cervical stroma has been described during late gestation and proposed as playing an essential role in the remodeling of cervical stroma tissue, essential for cervical ripening in preparation for parturition (40). The cervix is referred to as the “gatekeeper” of pregnancy and as such, a premature recruitment of immune cells into the cervix in response to lower reproductive tract infection has been proposed to induce premature loss of cervical integrity playing a key role in pre-term birth. Cervical macrophage infiltration is well reported in pre-term and term cervix in human and animal models (41). Abortion and preterm birth are well described in response to ZIKV in humans and NHPs including baboons, as we have described (25). Macrophages can induce cervical connective tissue remodeling via their expression of matrix metalloproteinases (MMPs), and various other factors that help in the breakdown of collagen and junction proteins resulting in the loss of cervical epithelial integrity required for cervical ripening (42). In relation to this, we observed ZIKV IF in the cervix of one FP isolate inoculated dam and all three PR isolate inoculated dams (Fig 3). For both isolates, the ZIKV IF was localized to the epithelial layer of the cervix in both endo- and ectocervical regions. The strongest ZIKV IF intensity was localized at the basal aspect of the epithelial cells at the junction with the stromal layer, which consists primarily of fibroblasts and smooth muscle cells. Macrophage IF was observed in both the FP isolate dam exhibiting ZIKV IF in cervical epithelium as well as all three ZIKV IF positive PR isolate dams in the epithelium (Fig 4). Relative to this, macrophages were routinely observed in the cervix of only the FP isolate dam with cervical ZIKV IF and the three PR isolate dams. In the FP dam, macrophages were observed in the stromal tissue immediately adjacent to the epithelium, indicative of recruitment (of monocytes) in response to the virus itself or from a local inflammatory reaction in response to ZIKV infection of epithelial cells. In the PR infected dams, macrophages were also noted adjacent to the epithelial layer as well as deeper in the stromal tissue, possibly indicating potential breakdown of the epithelial-stromal barrier and entry of virus into the stromal tissue. In contrast, FP dams with no ZIKV IF staining in cervix had only occasional macrophage staining in the stromal layer, typically scattered throughout the stroma. It is possible that the recruitment of macrophages due to ZIKV infection of the cervix through vaginal route at mid-gestation may induce breakdown of epithelial cell barrier and integrity similar to during cervical ripening at term. This breakdown could potentially lead to viral access to the adjacent reproductive tissues such as the uterus and placenta but more importantly, fetal membranes which lie at the top of the cervical canal thus amniotic sac and fluid and ultimately, the fetal compartment, thus exposing the fetus to Zika viral infection.

Vertical transfer of ZIKV in macaques appear to be very efficient, described to occur at near 100% following infection using various isolates of ZIKV including FP (37, 38), PR (31, 35, 43), BR (32, 44) and RIO (45, 46) isolates. While we observed no placental infection in the FP isolate inoculated pregnancies, vertical transfer to the placenta was observed (both RNA and IF) in two of the three animals infected with the PR isolate (Fig 5). In these two animals (dams PR1, 2), ZIKV RNA was detected in multiple cotyledons indicating widespread targeting of the placenta. Similar to our prior study infecting dams with the FP isolate (25), ZIKV IF was noted in the trophoblast cells. Since the one PR isolate inoculated dam without placental ZIKV targeting also was the dam with no noted viremia and latent detection in the CVW, this animal may have been infected by a later inoculation while it is clear that dams PR1 and 2 were infected at the first inoculation based on viremia, exhibited ZIKV RNA in the uterus and had prolonged ZIKV in the CVW. These dams also had notable ZIKV IF in the fetal membranes indicating that breakthrough of the virus through the placental barrier may have occurred since the membranes used for IF were at the uterus-placental interface or possibly via loss of cervical integrity leading to the opening of cervical canal and access to the membranes. While we found ZIKV RNA and IF in the placenta of two PR inoculated dams, there was no evidence of vertical transfer of the virus to any of the fetal compartments in any of the animals inoculated. The route of ZIKV infection, subcutaneous vs. vaginal may affect the rate and frequency of vertical transfer to the fetus per se. Considering the prolonged presence of virus in the CVW, the presence of ZIKV RNA and IF in placental trophoblasts and fetal membranes, and the unanticipated re-emergence of viremia (prolonged viremia) in these two PR inoculated dams, we predicted that vertical transfer would have occurred in these baboons at a later period, Further studies are needed to follow intravaginally ZIKV infected pregnant baboons for longer periods post-infection to better understand the fetal outcome of delayed viremia and potential re-emergence from immune privileged sites harboring ZIKV such as the lymph nodes.

With regard to the adaptive immune response to ZIKV infection, all six of the baboons inoculated with ZIKV developed ZIKV-specific IgM and IgG responses (Fig 2). IgM production following ZIKV infection was noted in all animals at variable times which indicated that the maternal immune system had access to the virus despite the lack of viremia in one of the three animals inoculated with the PR isolate. While, IgG titers were detected in all the dams 21 dpi, this was either too slow or inadequate to prevent spread of the virus to various reproductive tissues. It is noteworthy that the IgM and IgG response was also delayed in the dam inoculated with the FP isolate that displayed delayed viremia (11 dpi; FP3), being observed initially at 21 days (IgM) and 35 dpi (IgG).

Also, noteworthy, the one PR isolate infected dam that did not exhibit viremia had a similar weak IgM response (21 dpi) as well as IgG (35 dpi), with the immune cells likely being exposed to ZIKV via lymph nodes. It is unclear if these dams would have developed a robust neutralizing IgG response if examined at later times post-inoculation. As such, we can only speculate that some instances of sexual transmission of ZIKV may not result in a robust, neutralizing adaptive immune response.

While we acknowledge the small animal numbers in our study comparing PR and FP ZIKV isolates, there are clear indications that the PR isolate was more virulent in comparison to the FP isolate in terms of level of viremia, re-emergence of viremia, targeting of reproductive tissues and importantly, infection of the placenta and high potential for vertical transfer to the fetus per se. Few studies to date have focused on the effect of accumulated mutations in the virus acquired in the Americas compared to either the ancestral Asian isolate or the French Polynesian isolate that acquired the noted S139N substitution in the prM protein. Brazilian isolates, most notably the Rio isolate (RIO-U1/2016) acquired additional substitutions and the PR isolate differs in several other residues, some common with the Rio isolate (Fig 8). How these mutations may have increased virulence remains equivocal. However, viral modulation of the host immune response is a necessary factor for infection of the host and propagation of the progeny virus. All viruses must encode for at least one protein in their genome to modulate the host response to establish a successful infection. In the case of flaviviruses, many of the nonstructural proteins interact with cellular signaling cascades to instate a favorable environment for viral replication through resistance of host defense mechanisms (47). For flaviviruses, host response modulation is focused on the interferon response, and the level of modulation is directly proportional to pathogenicity as well as host species specificity (48). The increased virulence noted here for the Puerto Rican Strain (PRVABC59) compared to the isolate from French Polynesia (H/FP/2013) could be interpreted as a difference in modulation of host response. Alignment of the complete polyprotein from H/PF/2013, Brazilian RIO isolate, and PRVABC59, revealed specific amino acid changes in non-structural proteins NS1 (Lys1059→Glu) and NS5 (Ala2611→Val) of PRVABC59 that were not present in either H/PF/2013 or the RIO isolate (Fig 8). NS1 and NS5 play important roles in flavivirus replication, but have also been implicated in modulation of the host response through interaction with a variety of host proteins (49–51). Targeting of the interferon response by NS1 and NS5 has been documented for multiple flaviviruses including ZIKV (52–54) and the inhibition of IFNβ by NS1 can be mapped to a specific amino acid residue (49). Considering this, along with the multiple and complex ways flavivirus non-structural proteins manipulate the host response, the single amino acid changes identified in NS1 and NS5 for PRVABC59 could account for the increase in virulence for this specific isolate through altered interactions with host cellular proteins. However, further investigation is necessary to confirm this hypothesis. Domain III of the flavivirus envelope protein participates in receptor recognition and contains linear epitopes recognized by neutralizing antibodies (55). Interestingly, the single amino acid change in the envelope protein of PRVABC59 resides in domain III (Val620→Leu, Fig 8). This could possibly be due to the selective pressure of neutralizing antibodies as the virus moved from French Polynesia through Brazil into Puerto Rico. However, the two residues are highly similar hydrophobic amino acids, and it is highly likely the change is of no consequence to viral pathogenicity.

In conclusion, this study further clarified the transmission of ZIKV following intravaginal inoculation during pregnancy in a novel non-human primate model. This important translational model not only more closely recapitulates the course of observed infection patterns in humans, it also offers a novel comparator of the infectivity of two contemporary ZIKV isolates. The FP isolate, which was more rarely associated with vertical transmission than the PR isolate, appears to follow the same pattern within this study, which is logical given that ZIKV has continued to mutate during its passage from French Polynesia to the Americas. Future studies, some of which our lab is currently undertaking, can focus on the long-term outcomes of ZIKV viral infection following vaginal inoculation during early and mid-gestation in pregnancies that are allowed to continue gestating until term. This would be of value as sexual transmission of flaviviruses such as ZIKV may allow viral persistence or transmission to geographic locales when mosquito transmission is less likely, such as in the winter season. It is also a potential mechanism by which ZIKV infection can be spread to a population naïve to the ZIKV infection. Additionally, the course of ZIKV infection following sexual transmission and its consequences to the fetus appear different from subcutaneous ZIKV infection and what that means for the developing fetus and vaccine development is yet to be elucidated. This knowledge may help develop guidelines, preventative measures and therapeutics to protect against sexual transmission of ZIKV.

## ACKNOWLEDGEMENTS

Oklahoma Baboon Research Resource; Helen Lazear, University of North Carolina Chapel Hill, Mukesh Kumar, Georgia State University, Atlanta, GA, for providing ZIKV stocks; Oklahoma medical research foundation (OMRF) for tissue processing and immunostaining and James Tomasek, Vice President for Research, University of Oklahoma Health Sciences Center.

